# Sharper, Straighter, Stiffer, Stronger: Bill shape of male Green Hermits (*Phaethornis guy*) confers superior biomechanical performance during stabbing simulations

**DOI:** 10.1101/2025.03.02.641075

**Authors:** Felipe Garzón-Agudelo, Lucas Mansfield, Kevin Epperly, Alejandro Rico-Guevara

## Abstract

In hummingbirds, bill sexual dimorphism has been mainly related to differential use of floral resources between the sexes (i.e., intersexual resource partitioning). However, intrasexual selection has a potential role in driving hermit bill morphology. Males of *Phaethornis longirostris* possess weaponized bills, sharp and elongated dagger-like bill-tips, that enhance puncturing ability and territory defense during male-to-male combat at leks. In this study, we employed 3D modelling and finite element analysis to explore bill dimorphism and biomechanical stabbing performance in *Phaethornis guy.* We found that *P. guy* also exhibit a dimorphic weapon, with males displaying significantly sharper bill-tips than females. Additionally, we demonstrated a greater degree in biomechanical performance during horizontal stabbing in the straighter bills of male *P. guy* through a reduction in the energy expended in deformation (strain energy) and the risk of breakage (von Mises stress). Our findings indicate another example of bill-tip weapons and support the potential role of sexual selection in the evolution of hummingbird bill dimorphism.

## Introduction

Sexually dimorphic traits, beyond those directly related to sexual reproduction (e.g., reproductive organs), are a widespread consequence of the life histories of organisms across the animal kingdom (Janicke and Fromonteil, 2021; Rico-Guevara and Hurme, 2019). These intraspecific sexual differences can be expressed through distinct coloration, body size, or ornamentation (Andersson, 1994). Ecological mechanisms such as competition for resources (Butler et al., 2000) and differences in response to environmental gradients (Hendry et al., 2006) have been demonstrated to influence phenotypic differences between sexes. The evolutionary pressures behind sexual selection, a process wherein individuals compete with members of the same sex for access to reproductive opportunities, have also been linked to the development of secondary sexual dimorphism (Emlen, 2008; see Rico-Guevara and Hurme, 2019 for a thorough discussion of the hypotheses regarding the evolution of secondary sexual traits). Elucidating the specific mechanisms that drive sexual dimorphism within a species is a crucial step in deciphering its evolutionary history.

Hummingbirds make particularly excellent subjects for studying sexual dimorphism and its causative mechanisms, as they often exhibit notable intraspecific sexual differences (e.g., in plumage: Bleiweiss, 1997, Beltrán et al. 2022). Bill sexual dimorphism is a widespread pattern across the Trochilidae family (Berns and Adams, 2013), with females typically having longer bills than males, except in many species of the hermit clade (Phaethornithinae), wherein males tend to be the longer-billed sex (Bleiweiss, 1999, Temeles et al., 2010). Hermits also frequently exhibit strong sexual dimorphism in bill curvature, particularly in larger species, wherein females bear curvier bills than their male counterparts (Temeles et al., 2010). These differences have been primarily attributed to ecological causes related to feeding, wherein bill curvature allows each sex to exploit different floral resources (i.e., Intersexual resource partitioning: Temeles et al., 2010), with longer bills enabling one sex to access a broader range of flowers (Bleiweiss, 1999).

Hermits form leks during the breeding season, where males engage in aggressive behaviors such as vocalizations, visual displays, chases, and fighting (Snow, 1973; Stiles and Wolf, 1979; Harger and Lyon, 1980, MacDougall-Shackleton and Harbison, 1998). In Long-billed Hermits (*Phaethornis longirostris*), documented combat includes hovering males charging perched rivals and stabbing them in the throat with their bill-tip, followed by a chase and the intruder’s retreat (Movie A2 from Rico-Guevara and Araya-Salas, 2015). Adult males of *P. longirostris* bear a dagger-like structure present on the bill-tips, which is absent or reduced in females and juveniles. This dagger-like structure, characterized by a sharp and elongated maxillary tip, confers adult males a higher puncturing ability, and consequently higher success in defending lek territories (Rico-Guevara and Araya-Salas, 2015). It is possible that during stabbing, the straighter bills of males transmit more force without bending, and their pointer bills better transform that force into puncturing than blunter bills (Rico-Guevara and Araya-Salas, 2015, Rico-Guevara et al., 2019)—however, no studies have been conducted on the biomechanical demands (e.g., strain energy) experienced by hummingbird bills during stabbing. The puncturing ability of male bills and its relationship with lekking behavior suggest that sexual dimorphism in the bills of *P. longirostris* is caused by intrasexual selection, promoting more weaponized bills (Rico-Guevara and Araya-Salas, 2015). This idea provides a complementary hypothesis to the notion that sexual dimorphism is the result of intersexual resource partitioning (Temeles et al., 2010), and is a seemingly rare example of intrasexually selected weaponry in birds (Rico-Guevara and Hurme, 2019).

To further explore the weaponization hypothesis for bill sexual dimorphism in hermits and determine if weaponization is unique to *P. longirostris*, we chose to study the Green Hermit (*Phaethornis guy*), as it is a large species that displays fighting lekking behavior (Snow, 1973) and bill sexual dimorphism (i.e., longer and straighter bills in males; Temeles et al., 2010). Additionally, *P. guy* is closely phylogenetically related to *P. longirostris* (McGuire et al., 2014), a known bearer of a weaponized bill (Rico-Guevara and Araya-Salas, 2015). We explored bill sexual dimorphism and its relationship to stabbing performance using 3D models and finite element analysis simulations. We expected *P. guy* to show intersexual morphological differences in bill-tip sharpness, indicative of the presence of a dagger-like trait in males as seen in *P. longirostris*. We hypothesized that the straighter bills of *P. guy* males perform better as stabbing weapons relative to females. We expected male bills to be structurally stronger (lower stress), more efficient at transmitting forces (higher stiffness: lower strain energy), and more resistant to buckling (higher buckling load factors) under the application of axial loads.

## Materials and Methods

### Specimen information and photogrammetry modeling

We selected eight female and eight male *Phaethornis guy* specimens, housed at the Burke Museum’s ornithology collection (see specimen IDs in Table S1). All selected specimens had well-preserved, undamaged bills, and were classified as adults based on the low percentage of maxilla covered by corrugations (Yanega et al., 1997), the relative gonad size (Stiles, 1980), and the absence of Bursa of Fabricius (Rhodes et al., 2022). The male specimens exhibited two plumage varieties, being either gray-chested or green-chested individuals. Prior research indicates that the gray-chested plumage is lost over the course of four years, indicating that our gray-chested birds are younger adult males (Snow, 1974; Stiles and Wolf, 1979). However, we did not consider male color variation in our analyses due to the limited sample size. Three-dimensional (3D) models of the bills were generated using high-resolution 3D digital photogrammetry following the methods of Medina et al. (2024). Briefly, for photogrammetry data collection, we used six Sony ⍺7R mirrorless cameras, each equipped with Sony 90 mm macro lenses and synchronized via remote trigger (Vello, New York City, United States). To avoid potential measurement error due to differential separation of the jaws across individuals, likely introduced as a specimen preparation artifact, we used only the maxilla in our analyses (hereafter, *bill* refers to the maxilla).

### Bill sexual dimorphism

We evaluated bill sexual dimorphism using landmark-based geometric morphometrics and by measuring bill-tip sharpness, bill curvature, arc length, and surface area (Fig. 1A). Three specimens were excluded from bill shape analysis and surface area calculations due to peeling in the keratin of the rhamphotheca and lateral shifts in the fit of the maxilla and mandible, causing the mandible to slightly cover one of the tomia, which would affect measurements of surface area and shape analysis. Ensuring a natural alignment between upper and lower bills during specimen preparation is crucial to avoid measurement errors in morphometric analysis, particularly in small species where subtle shifts may go unnoticed.

**Fig. 1.**
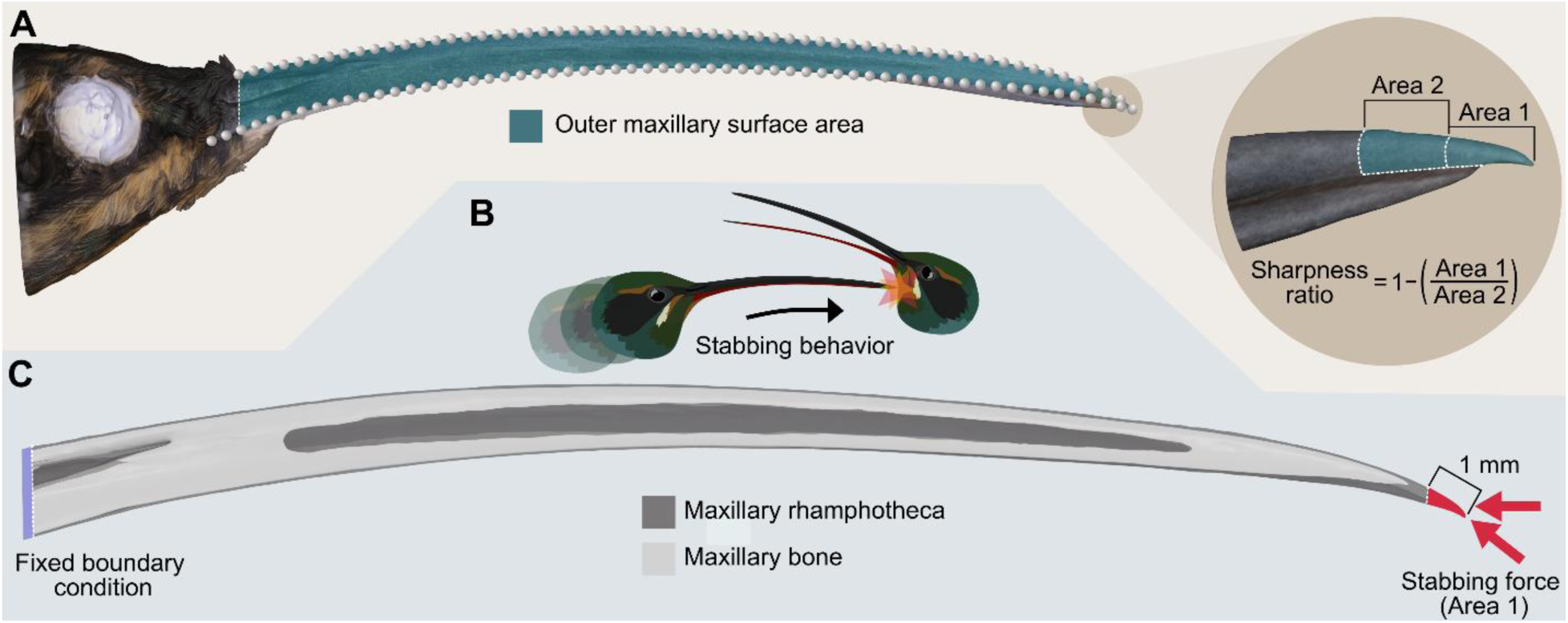
Morphological characterization of the bill and stabbing behavior simulation. (A) Lateral view of a *Phaethornis guy* 3D photogrammetry model (left), showing landmarks used to quantify bill shape variation (grey spheres) and selected area to quantify bill surface area (green shaded area). The landmarks on the culmen were used to measure bill arc length and bill curvature (arc:chord ratio). Close-up of the bill-tip (right) showing the outer maxillary areas used to calculate the sharpness ratio. Area 1 captures the area of the distal-most millimeter of the bill, and Area 2 of the second-most distal millimeter. (B) *P. guy* hypothetical stabbing behavior based on *P. longirostris* fight (Movie A2) from Rico-Guevara and Araya-Salas (2015). (C) Boundary conditions for finite element analyses simulating stabbing behavior with 3D CT models. Loads were applied at two different angles from the bill-tip.

Prior to all measurements, the 3D photogrammetry models were imported to Blender (Blender Foundation; www.blender.org), where the bill longitudinal axis was aligned parallel to the X-axis. We then used 3D Slicer (Fedorov et al., 2012) to digitize three contour lines on the bill models, one traced from the bill-tip along the medial-dorsal maxillary surface (in cross section, the line would go through the tallest point of the bill) to the proximal end of the exposed culmen (where the keratinous surface of the bill meets the feather line), and two other lines along the left and right tomia, from the bill-tip to the most proximal points of the tomia. Each contour curve was resampled to 60 points and intersecting points at the bill-tip were removed, giving a total of 178 points (Fig. 1A). The endpoints of the curves were treated as homologous landmarks (4 points) and the rest as semilandmarks (174 points; Gunz and Mitteroecker, 2013). The raw landmark coordinates were imported to R for analysis. We removed non-shape variation (*i.e.*, position, scale, and orientation) through a Generalized Procrustes Analysis (GPA; Rohlf, 1999) using the “gpagen” function in the Geomorph R package (Baken et al., 2021). Semilandmarks were allowed to slide using the Procrustes distance minimization method, following the recommendations of Perez et al. (2006). As bills are bilaterally symmetric structures, the symmetric component of shape variation was extracted with the function “bilat.symmetry” and used to obtain a representation of the bill shape space through a Principal Component Analysis (PCA) with the function “gm.prcomp” from Geomorph. As a result of GPA, shape (Procrustes coordinates) and size (centroid size) variables were obtained.

We used the landmark coordinates of the culmen to measure bill curvature and arc length (Fig. 1A). We treated curves as two-dimensional and measured curve dimensions with the “coo_scalars” function in the Momocs R package (Bonhomme et al., 2014). Bill curvature was calculated using the arc:chord ratio (Stiles, 1995), where arc refers to the length of the upper curve of the bill from the bill-tip to the proximal point of the exposed culmen (arc length), and chord to the linear distance between these two points. Similarly to Rico-Guevara and Araya-Salas (2015), we used arc:chord ratio as it was the most conservative method for bill curvature calculation from Berns and Adams (2010), to ensure that any measured differences were of biological importance.

To quantify surface area and sharpness (Fig. 1A), the models and their textures were imported into MeshLab (Callieri et al., 2012). Uploading the textures allowed us to clearly visualize the tomia (cutting edges of the beak). We used the “subdivision surface tool” to refine the mesh at the tomia and the bill-tip without altering the model shape, the “face selector tool” to select the outer area of the maxilla from the bill-tip to the proximal end of the exposed culmen (in order to avoid differential covering of the maxilla by the face feathers at the basal end of the bill), and applied the “compute area of selection” function to extract the surface area.

The dagger of *P. longirostris* maxillary tip is characterized by being sharper and more elongated over the mandible in adult males (Rico-Guevara and Araya-Salas, 2015). Maxillary elongation cannot be reliably measured in museum specimens, as the fit between the maxilla and mandible shifts from the natural configuration with desiccation and specimen preparation artifacts. To measure bill-tip sharpness, we used the 3D equivalent of the “pointiness index” employed by Rico-Guevara and Araya-Salas (2015). We measured a 1mm distance from the tip of the bill and calculated the surface area of this distal area. We then measured a further 1mm towards the head and calculated the surface area for this basal segment. The distal measurement was divided by the basal segment and subtracted from 1 to produce a sharpness ratio, with greater values indicating sharper bills (Fig. 1A).

### CT modeling and finite element analysis

To generate 3D models containing information of the inner bill structure, we CT-scanned one female and one male located at the extremes of PC1 of the bill shape space (Fig. 2A, B, Table S1). The extremes of PC1 captured the most pronounced intersexual differences in bill shape observed in our sample (see Results and Discussion). Selecting these specimens allowed us to analyze the full range of morphological variation within our sample and its potential functional implications. Additionally, extreme morphologies are likely to reflect individuals with potentially enhanced adaptations to sexual selection pressures. Specifically, males at the extreme negative end of PC1 displayed straighter, sharper bills, which may confer an advantage in stabbing performance during intrasexual competition.

**Fig. 2.**
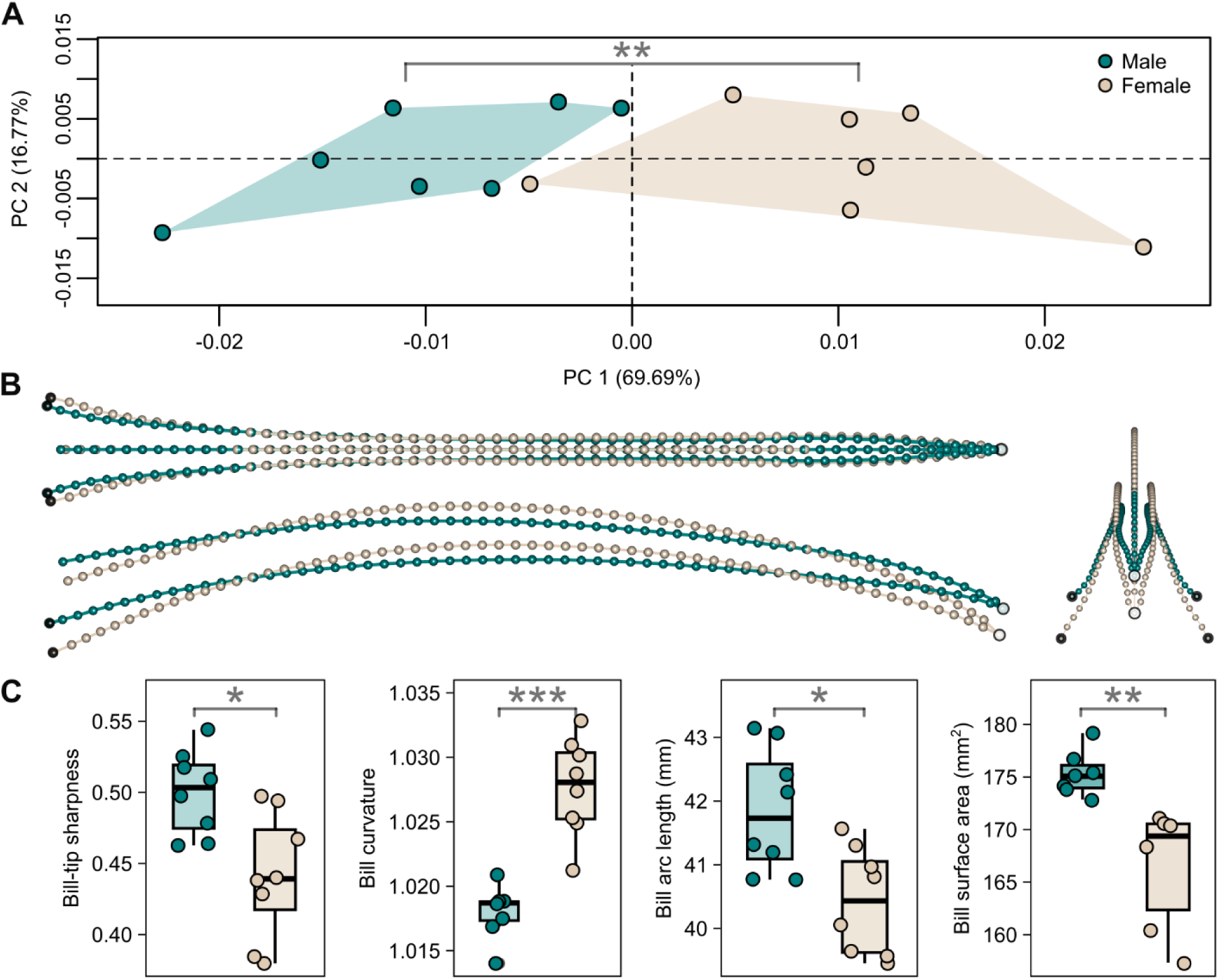
Bills of male *Phaethornis guy* were sharper, straighter, longer, and had greater surface area relative to females. (A) First 2 principal components of the symmetric component representing bill shape space. Plot size is scaled relative to the percentage of variation explained by PCs. (B) Bill shape changes associated with the extremes of PC1 in dorsal (top-left), lateral (bottom-left), and frontal (right) views. The green shape corresponds to the male with extreme bill shape (negative end of PC1), and the light brown shape to the female with extreme bill shape (positive end of PC1). Shape differences were not magnified. (C) Boxplots showing intersexual comparisons of bill-tip sharpness, bill curvature, arc length, and surface area. Level of significance: \**P*<0.05, \*\**P*<0.01, \*\*\**P<*0.001.

Scans were performed using an NSI X5000 scanner with a voltage of 100 kV, a current of 100 µA, and a voxel resolution of 27 µm. TIFF images from CT scans were imported to 3D Slicer (Fedorov et al., 2012) and used to create 3D models of the maxillary rhamphotheca and bone in STL (stereolithography) format. Using Geomagic Studio 2014 (Geomagic Inc., Research Triangle Park, NC, USA), we isolated the bill from the head at the proximal-most point of the exposed culmen, removed any model artifacts, aligned the bill-tip and the tomia proximal ends to a common horizontal plane, and converted the STL models into CAD (computer-aided design) geometries. The resulting watertight (*i.e.*, closed-surface) 3D models were imported into Ansys 2023 R2 (ANSYS Inc., Canonsburg, PA, USA) for linear static finite element analysis (FEA; Rayfield, 2007).

We then generated FE meshes from the CAD models, composed of quadratic (10-noded) tetrahedral elements (Female model: 878,768 nodes and 464,982 elements; Male model: 923,742 nodes and 492,350 elements). Homogeneous and isotropic material properties were assigned to the meshes, with Young’s moduli of 1700 MPa for the rhamphotheca and 7300 MPa for the inner bone, and a Poisson’s ratio of 0.4 for both tissues, based on parameters previously used for bird bills (Soons et al., 2012a, 2012b). Bonded contacts *(sensu* Marcé-Nogué, 2022) were used between the rhamphotheca and the bone. The models were constrained at the base of the bill to represent the area of attachment to the head. Axial forces were then applied to the last millimeter of the bill-tip to simulate stabbing combat behavior (Fig. 1B, C) as seen in *P. longirostris* (Movie 2 from Rico-Guevara and Araya-Salas, 2015). To assess performance under different conditions, forces were applied at two angles: one horizontally (horizontal stabbing) and another parallel to the axis of the bill-tip (parallel stabbing).

We conducted two types of FE analyses: one in which we varied the magnitude of the applied forces based on empirical data (unscaled models), and another where forces were scaled according to equivalent force-to-surface-area ratios (scaled models). For unscaled analyses, we used the average forces required by the bills of *P. longirostris* to pierce a polyvinyl chloride film—200 mN for females, and 125 mN for adult males—as previously measured by Rico-Guevara and Ayalas-Salas (2015). The unscaled models allowed us to make intersexual comparisons of bill mechanical behavior while performing the hypothesized function of dagger-like bill-tips (i.e., puncturing). Scaled models, on the other hand, allow the comparison of mechanical performance due to shape alone by removing size differences (Dumont et al., 2009). For scaled analyses, we removed differences in size by choosing a reference model (A) and scaling the loads applied to the target model (B) to an identical force (F): surface area (SA) ratio using Eqn 1 (Dumont et al., 2009). For this, we used the surface area of the entire CT model (Female SA: 322.934 mm^2^; Male SA: 327.791 mm^2^). We selected the female specimen as the reference model (Fig. 2A, Table S1). It is important to note that the choice of reference model in FE comparative analysis is arbitrary. The resulting scaled force value applied to the male model was 203.01 mN.

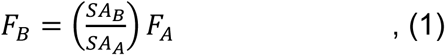

We used three metrics to compare the relative stabbing performance of *P. guy* bills: von Mises (VM) stress, total strain energy, and buckling load factor. VM stress is a good predictor of failure under ductile fracture, so, for a given load, structures with lower VM stress are less likely to fail, indicating that they are structurally stronger (Dumont et al., 2009). Total strain energy is a measure of the work expended in deformation, with lower values indicating stiffer structures that transmit force more efficiently (Dumont et al., 2009). Since long, slender structures subjected to axial loads may fail through buckling (*i.e.*, sudden change in shape due to loss of stability under compressive loads; Wainwright et al., 1976; Marcé-Nogué, 2022), we assessed buckling resistance using eigenvalue buckling analysis (Cook et al., 2002), wherein the minimum eigenvalue corresponds to the buckling load factor. Higher buckling load factor values indicate greater resistance to buckling failure. We note that these values should only be considered in a comparative context; they do not represent absolute measures of bill mechanical performance.

### Data analysis

To assess sexual dimorphism in bill shape of *P. guy*, we performed Procrustes ANOVA with permutation procedures (Goodall, 1991) using the “procD.lm” function from Geomorph. With the same function, we explored the relationship between bill curvature and sharpness to bill shape, and between bill length and surface area to bill centroid size. We did not test for allometric effects, as all specimens were adults. We used non-parametric Mann-Whitney U Tests to evaluate intersexual differences in the bill-tip sharpness, bill curvature, arc length, and surface area. All calculations, as well as morphometric and statistical analyses, were performed in R version 4.4.1 (R Core Team, 2024; http://www.R-project.org/).

We evaluated relative bill stabbing performance by comparing the magnitudes of VM stress, total strain energy, and buckling load factor between the sexes. These single-value metrics are useful indicators of the biomechanical performance of a model. We summarized VM stress using the peak stress and the mesh weighted arithmetic mean of the stress (MWAM). Artificially high stress values, known as numerical singularities, frequently occur in FE models of complex shapes, potentially obscuring the identification of biologically relevant results (Marcé-Nogué, 2020). We excluded the top 2% stress values to avoid these artifacts, using the 98th percentile as the peak VM stress—an approach applied by previous studies (Dumont et al., 2014; Habegger et al., 2019; Sansalone et al., 2024). Instead of the mean stress, we used MWAM because it accounts for differences in element volumes within the meshes (Marcé-Nogué et al., 2016; Murphy et al., 2024; Rowe and Rayfield, 2022). The distribution of VM stress was visualized using a variation of the inferno color map (*i.e*., plasma), as it offers increased accessibility for people with color vision deficiency and reduced distortion of the data (Lautenschlager, 2021).

## Results and discussion

### Bill sexual dimorphism

Our bill-tip sharpness index indicates that *P. guy* males possess dagger-like bill-tips, candidate traits to be considered intrasexually selected weapons (Rico-Guevara and Hurme, 2019), as they are present or more developed in one sex, but their use as contact-contest tools during same-sex agonistic encounters warrants further field-based investigation. However, given that lekking species of *Phaethornis* exhibit similar aggressive behaviors (Snow, 1974; Stiles and Wolf, 1979), and the congeneric *P. longirostris* does use its bill as a weapon (Rico-Guevara and Ayalas-Salas, 2015), we expect that *P. guy* uses it as well. Whether *P. guy* daggers enhance reproductive success through lekking territory tenure remains to be explored.

We found significant intersexual differences in both the shape and size of *P. guy* bills. Sexual dimorphism aligned closely with PC1 of the bill shape space (Fig. 2A), showing a clear separation between sexes along PC1 (69.69%) and high overlap along PC2 (16.76% of total shape variation). Intersexual differences accounted for 45% of bill shape variation (*R^2^* = 0.45, *P*= 0.004). Curvature contributed substantially to bill shape variation (*R^2^* = 0.57, *P*= 0.001), and was related to shape differences observed along PC1. Namely, male bills towards the negative side of PC1 were straighter, and female bills towards the positive side were curvier (lateral view in Fig. 2B). Males had significantly straighter (*W*=64, *P*<0.001) and longer bills than females (*W*=11, *P*=0.03; Fig. 2C), as previously seen in *P. guy* and other hermit species (Bleiweiss, 1999; Temeles et al., 2010; Berns and Adams, 2013). Males had significantly sharper bill-tips (*W*=8, *P*=0.01; frontal view in Fig. 2B; Fig. 2C) and greater bill surface area (*W*=0, *P*=0.0012). Sharpness explained 13% of bill shape variation, but this relation was not significant (*R^2^* = 0.13, *P*= 0.161). Bill centroid size had a stronger association with bill arc length (*R^2^* = 0.94, *P*= 0.001), than with surface area (*R^2^* = 0.69, *P*= 0.001).

Our findings align with previous research on bill curvature and length dimorphism in hummingbirds, which have been linked to ecological causes. In species-poor environments, the absence of competing species enables males and females to occupy separate ecological niches that would otherwise be filled by closely related species, with bills of distinct curvature allowing both sexes to specialize on feeding from different flowers (Temeles et al., 2010). Longer bills allow one sex to access a greater variety of floral resources, while shorter, straighter bills enhance efficiency in feeding from small, localized patches (Bleiweiss, 1999). However, sexual selection could also drive similar and additional bill features by enhancing combat effectiveness during lekking territory defense. Longer bills may increase reach during male-male confrontations, while sharper bill-tips would increase puncturing inflicted on the opponent (Rico-Guevara and Ayalas-Salas, 2015, Rico-Guevara et al. 2019). Furthermore, *ceteris paribus*, an increase in surface area leads to a larger cross-sectional area, which in turn enhances bending strength (Wainwright et al., 1976).

### Bill stabbing performance

Our results demonstrate that the straighter shape of male *P. guy* bills provides biomechanical advantages for horizontal stabbing, primarily by efficiently transmitting fighting forces. In the scaled analysis, the male bill expended 51.2% less energy in deformation, the largest difference observed across all metrics (Fig. 3A), indicating that male bills are more efficient at transmitting stabbing forces at horizontal angles than female bills. Furthermore, the male showed a lower risk of breakage, with a 39% reduction in MWAM and a 17.5% reduction in peak von Mises stress relative to the female. In contrast, when stabbing at the angle parallel to the bill-tip, the biomechanical performance was more similar between the sexes, with differences in von Mises stress and total strain energy remaining modest (<10%, Fig. 3B). Both sexes exhibited comparable buckling resistance under both loading angles (around 10%, Fig. 3). Relative to the female, the male did not experience a substantial loss of stabbing performance across tested angles. Instead, its performance increased under horizontal loading, suggesting that the straighter bills of males allow for a greater range of potential stabbing angles, requiring less precision during fighting contests.

**Fig. 3.**
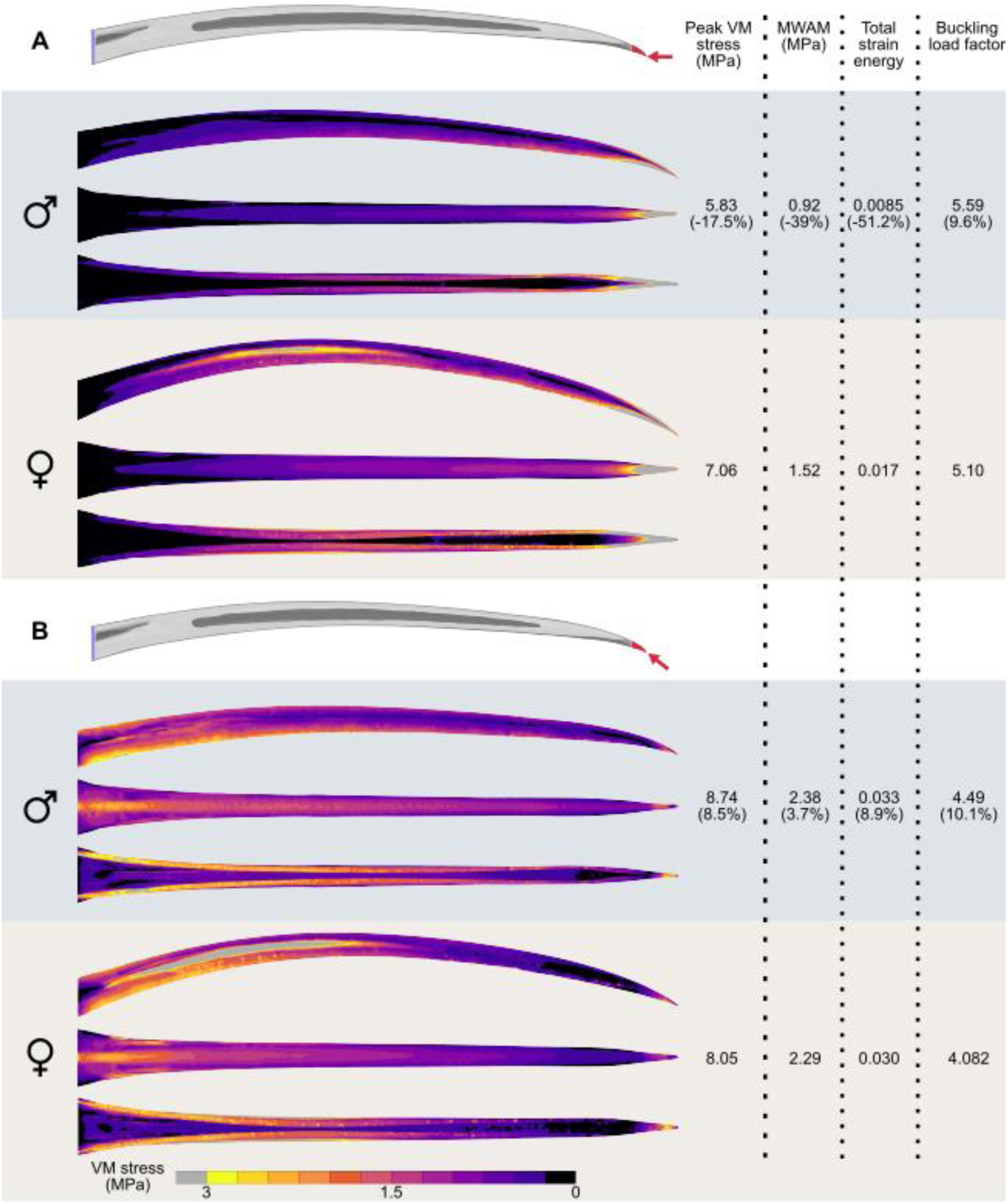
Relative to females, bills of male *Phaethornis guy* were stronger and stiffer under horizontal scaled loading. Scaled FEA results of stabbing simulation at two load angles: (A) horizontal and (B) parallel to the bill-tip axis. Left: Contour plots showing von Mises stress distribution on male and female bills in lateral, dorsal and ventral views. Warmer colors indicate regions of higher stress. Right: Comparisons of biomechanical performance metrics including peak and mesh weighted arithmetic mean von Mises stress, total strain energy, and buckling load factor. Values in parenthesis represent the percentage difference of male values relative to the reference female.

In the unscaled analyses, the male bill showed considerably greater biomechanical performance in both stabbing scenarios (in all metrics, Fig. S1), with larger intersexual differences in strength and stiffness observed during horizontal loading. Lower forces were applied to the male bill in the unscaled compared to the scaled analyses, reflecting that sharper tools can penetrate target materials with lower forces due to the reduced area over which the force is concentrated (Anderson, 2018). Rico-Guevara and Ayalas-Salas (2015) demonstrated that the sharper bills of male *P. longirostris* require less force to pierce a skin-like material compared to females. Thanks to their sharper bill-tips, male *P. guy* may minimize the forces used to accomplish puncture, subjecting their bills to lower biomechanical demands. Direct measurements of stabbing forces during lekking interactions would provide valuable insights into the actual biomechanical demands experienced by bills in combat.

Across both scaled and unscaled models, parallel loading consistently produced higher stress and strain energy values, and lower buckling load factor values than horizontal loading. The stress propagation patterns differed between the two loading angles. During horizontal loading, bills experienced downward bending, with regions of high von Mises stress concentrated at the bill-tip and distributed along the tomium and culmen, dissipating longitudinally toward the base of the maxilla. In parallel loading, bills exhibited upward bending, with high von Mises stress concentrated at the base and dissipating toward the tip, with a drop in stress around the distal quarter of the bill and a localized increase at the loading point. These differences in stress propagation indicate that bills experience distinct mechanical challenges depending on the stabbing angle, potentially influencing fighting strategies and angles of stabbing attacks. Additionally, under both loading conditions, the female bill showed a concentration of stress on the surface between the culmen and the tomium.

It is possible that *P. guy* could prefer to stab opponents at angles horizontal to the bill axis for different reasons. First, our results indicate that bills transmit forces more efficiently, experience less stress, and are more resistant to buckling during horizontal stabbing (Fig. 3; Fig. S1). In contrast, when loads were applied parallel to the bill-tip, bills exhibited lower strength and stiffness. This pattern aligns with findings for other curved piercing tools. Scorpion stingers and spider chelicerae have optimal piercing angles that minimize structural stress and strain energy (van der Meijden and Kleinteich, 2017; Bar-On et al., 2014). Notably, these optimal angles consistently differ from the angle parallel to the tip of the structure. Second, horizontal stabbing is a less complex behavior. It occurs when the attacker flies toward the target individual, and its bill-tip meets the target’s body. In comparison, parallel stabbing requires a more intricate motion, wherein the attacker must rotate its head ventrally following the curvature of the bill after approaching and contacting the target. It is important to note that the angle of attack during puncturing has little influence on the damage inflicted on the target (Zhang et al., 2024). Thus, hummingbirds may adjust their attack angles to minimize the stress and deformation experienced by their bills without compromising the inflicted damage. While our findings suggest a biomechanical advantage for horizontal stabbing, whether male *P. guy* prefer this angle during combat requires confirmation through behavioral studies at lekking sites.

### Concluding remarks

We show that male *P. guy* possess weaponized bills with sharp dagger-like tips, an intrasexually selected trait previously identified only in *P. longirostris* and one of the few known examples of male weaponry in birds (Rico-Guevara and Ayalas-Salas, 2015; Rico-Guevara and Hurme, 2019). Furthermore, our results demonstrate that the straighter shape of male bills provides biomechanical advantages for horizontal stabbing by efficiently transmitting fighting forces, supporting the potential role of sexual selection in the evolution of bill dimorphism in hummingbirds.

While 3D models have been widely used to study bill shape across bird species and higher clades (e.g., Cooney et al., 2017), they had not, to our knowledge, been used to examine bill sexual dimorphism. Likewise, FEA has been used in birds to evaluate biomechanical performance in food processing and cavity excavation (Attard et al., 2016; Chhaya et al., 2023), and in other taxa, such as arthropods and mammals, to study weapon function (McCullough et al., 2014; Klinkhamer et al., 2019), but this study represents the first application of 3D modeling and finite element analysis to address questions related to sexual selection in birds. The methods employed here could be part of a framework for studying the presence and function of bill weaponry across the avian class.

Although this study provides an accessible framework for bill weaponry research, our analysis nevertheless faced limitations. Aside from the specimens excluded as described in the methods, we used homogeneous material properties to isolate the effects of shape on biomechanical performance. However, bill elasticity may vary along its length, as suggested by flexural rigidity estimates in hummingbirds (Rico-Guevara et al., 2024). Additionally, male *P. longirostris* bill-tips feel stiffer to the touch (Rico-Guevara and Ayalas-Salas, 2015), possibly due to differences in mineralization. Stiffer bill-tips could enhance resistance to stabbing-induced stress but may reduce feeding efficiency, as maxillary-tip bending plays a key role in nectar extraction (Rico-Guevara et al., 2023, 2024). Studies on hummingbird bill material properties and the potential trade-off between stabbing performance and feeding efficiency are warranted.

Future work should explore bill-tip sharpening in other hummingbird species, both within hermits and across the Trochilidae family, examining its relation to lekking and feeding territoriality (Stiles and Wolf, 1979; Sargent et al., 2021), to better understand the distribution and evolutionary drivers of this trait. Detailed descriptions and quantification of hummingbird fighting behavior are needed. Of particular interest is the case of female hummingbirds in species with feeding territoriality, as their parental care duties may conflict with developing bills for stabbing. Sharp bill-tips could pose a risk of injury to nestlings during feeding (which involves deep insertion of the female’s bill-tips into the chicks’ throats), potentially limiting the evolution of this trait in females. If sharp bills do occur in female hummingbirds, it would be valuable to explore whether there are behavioral or morphological adaptations in females or nestlings to prevent injury. Weapons are considered rare in birds, as the high energetic cost of flight may prevent their evolution or promote their loss (Menezes and Palaoro, 2022). However, the presence of weaponry in hummingbirds suggests that such traits can evolve as fine modifications of preexisting structures like bills without increasing flight costs. We highlight the possibility that similar adaptations have evolved in other avian lineages that exhibit combat behaviors, potentially making weapons more widespread in birds than previously recognized.

## Acknowledgements

This work would not have been possible without the resources and support of the X-ray Computed Tomography Facility at the University of Washington, along with the Ornithology Department at the Burke Museum, which allowed us to CT scan several of their *Phaethornis guy* specimens. We are grateful to Madison Mayfield for her assistance in obtaining and processing CT scans of the specimens. We also thank the members of the Behavioral Ecophysics Lab at the University of Washington for their valuable insights and suggestions for improving this manuscript.

## Author contributions

Conceptualization: F.G-A., L.M., K.E., A.R-G.; Methodology: F.G-A., L.M.; Formal analysis: F.G-A., L.M.; Investigation: F.G-A., L.M.; Writing - original draft: F.G-A., L.M.; Writing - review & editing: F.G-A., L.M., K.E., A.R-G.; Visualization: F.G-A.; Resources: K.E., A.R-G.; Supervision: K.E., A.R-G.; Funding acquisition: A.R-G.

## Data availability

All relevant data are reported within the manuscript and supplementary information.

---

**Fig. S1.**
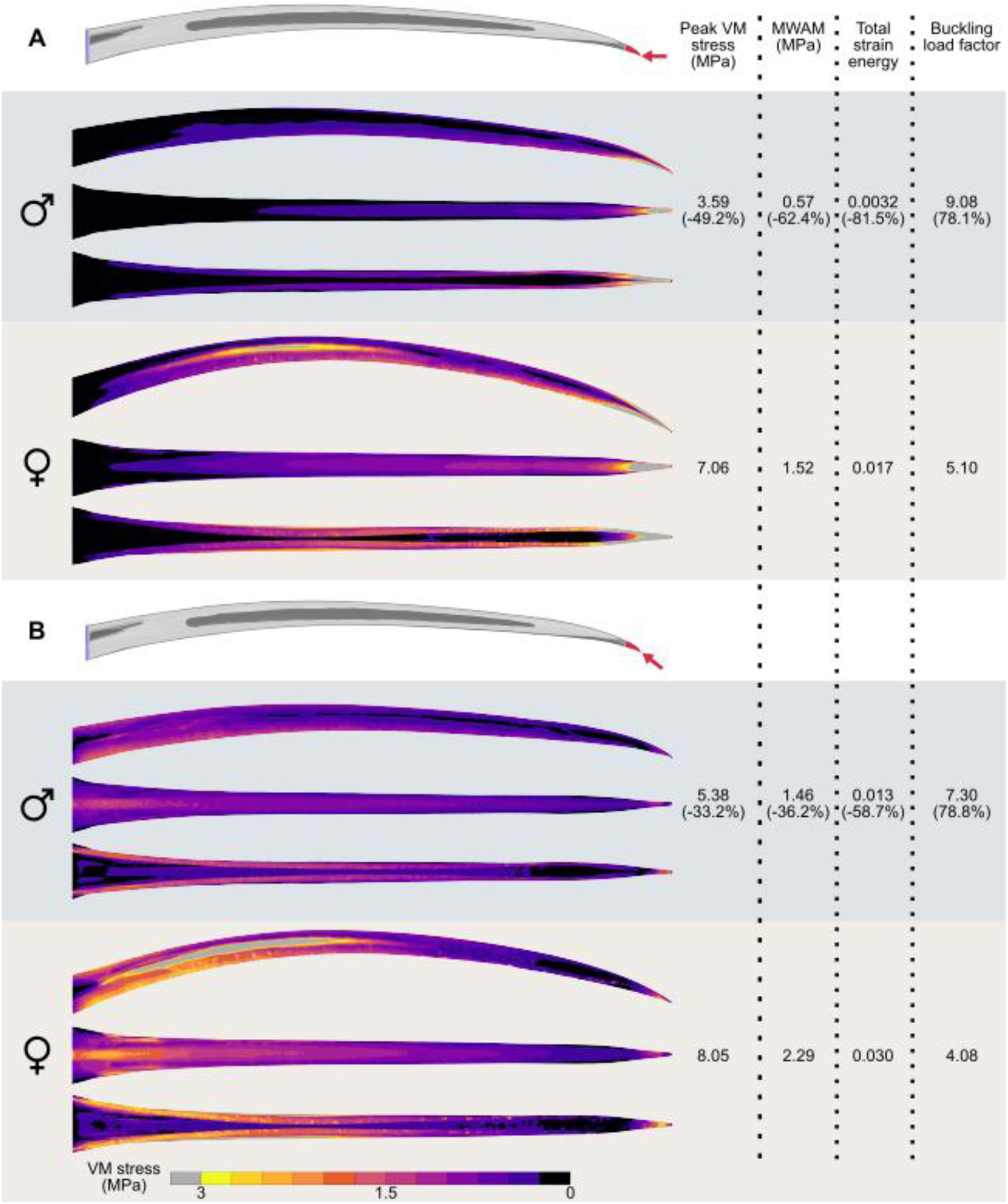
Relative to females, bills of male *Phaethornis guy* were stronger, stiffer and more resistant to buckling under horizontal and parallel *unscaled* loading. Unscaled FEA results of stabbing simulation at two load angles: (A) horizontal and (B) parallel to the bill-tip axis. Left: Contour plots showing von Mises stress distribution on male and female bills in lateral, dorsal and ventral views. Warmer colors indicate regions of higher stress. Right: Comparisons of biomechanical performance metrics including peak and mesh weighted arithmetic mean von Mises stress, total strain energy, and buckling load factor. Values in parenthesis represent the percentage difference of male values relative to the reference female. Under both loading scenarios, male bills showed greater resistance to breaking and to buckling failure, and expended less energy in deformation.

**Table S1.** Bill morphometric variables.

| Specimen ID | Sex | Sharpness | Arc:Chord | Arc length (mm) | SA (mm<sup>2</sup>) | CS (mm) | PC1 | PC2 |
| --- | --- | --- | --- | --- | --- | --- | --- | --- |
| UWBM-76887 | M | 0.525 | 1.019 | 40.767 | 172.847 | 160.177 | -0.0103 | -0.0035 |
| **UWBM-76890** | M | 0.544 | 1.017 | 41.195 | 174.079 | 161.784 | -0.0228 | -0.0093 |
| UWBM-76893 | F | 0.380 | 1.027 | 40.056 | 168.370 | 158.029 | 0.0113 | -0.0011 |
| UWBM-76984 | M | 0.463 | 1.014 | 43.068 | 173.824 | 166.616 | -0.0151 | -0.0002 |
| UWBM-108340 | M | 0.509 | 1.017 | 42.419 | 176.764 | 166.820 | -0.0068 | -0.0037 |
| UWBM-108783 | F | 0.384 | 1.021 | 41.305 | 171.049 | 162.935 | -0.0050 | -0.0031 |
| UWBM-111316 | F | 0.494 | 1.029 | 39.641 | 160.349 | 155.119 | 0.0048 | 0.0080 |
| UWBM-112274 | M | 0.517 | 1.019 | 42.141 | 179.167 | 165.861 | -0.0036 | 0.0071 |
| UWBM-112313 | F | 0.438 | 1.025 | 39.451 | 170.590 | 155.885 | 0.0105 | 0.0049 |
| UWBM-123358 | M | 0.478 | 1.019 | 43.144 | - | - | - | - |
| UWBM-123451 | M | 0.464 | 1.019 | 40.774 | 175.068 | 162.004 | -0.0116 | 0.0064 |
| UWBM-123474 | F | 0.497 | 1.025 | 39.568 | 157.342 | 155.097 | 0.0106 | -0.0065 |
| UWBM-123475 | F | 0.428 | 1.030 | 40.809 | - | - | - | - |
| UWBM-123546 | F | 0.440 | 1.033 | 41.565 | - | 163.425 | 0.0136 | 0.0056 |
| **UWBM-123686\*** | F | 0.467 | 1.031 | 40.968 | 170.380 | 162.295 | 0.0248 | -0.0111 |
| UWBM-125003 | M | 0.498 | 1.021 | 41.319 | 175.480 | 163.747 | -0.0005 | 0.0063 |

Selected specimens for FEA are in bold. The asterisk marks the reference specimen for scaled FEA. M, male; F, Female; SA, outer maxillary surface area; CS, centroid size; PC, principal component of shape variation.

## References

Andersson, M. (1994). Sexual Selection. Princeton University Press.

Anderson, P. S. L. (2018). Making a point: shared mechanics underlying the diversity of biological puncture. J. Exp. Biol. 221, jeb187294. 10.1242/jeb.187294

Attard, M.R.G., Wilson, L.A.B., Worthy, T.H., Scofield, P., Johnston, P., Parr, W.C.H., and Wroe, S. (2016). Moa diet fits the bill: virtual reconstruction incorporating mummified remains and prediction of biomechanical performance in avian giants. Proc. R. Soc. B. 283: 20152043. 10.1098/rspb.2015.2043

Baken, E. K., Collyer, M. L., Kaliontzopoulou, A., and Adams, D. C. (2021). geomorph v4.0 and gmShiny: Enhanced analytics and a new graphical interface for a comprehensive morphometric experience. Methods Ecol. Evol. 12(12), 2355–2363. 10.1111/2041-210X.13723

Bar-On, B., Barth, F. G., Fratzl, P., and Politi, Y. (2014). Multiscale structural gradients enhance the biomechanical functionality of the spider fang. Nat. Commun. 5(1), 3894. 10.1038/ncomms4894

Beltrán, D. F., Araya-Salas, M., Parra, J. L., Stiles, F. G., and Rico-Guevara, A. (2022). The evolution of sexually dimorphic traits in ecological gradients: an interplay between natural and sexual selection in hummingbirds. Proc. R. Soc. B. 289(1989). 10.1098/rspb.2022.1783

Berns, C. M., and Adams, D. C. (2010). Bill shape and sexual shape dimorphism between two species of temperate hummingbirds: Black-Chinned hummingbird (*Archilochus alexandri)* and Ruby-Throated hummingbird (*A. colubris)*. Auk. 127(3), 626–635. 10.1093/beheco/aru182

Berns, C.M., and Adams, D.C. (2013). Becoming Different But Staying Alike: Patterns of Sexual Size and Shape Dimorphism in Bills of Hummingbirds. Evol. Biol. 40, 246–260. 10.1007/s11692-012-9206-3

Bleiweiss, R. (1997). Covariation of sexual dichromatism and plumage colours in lekking and non-lekking birds: A comparative analysis. Evol. Ecol. 11, 217–235. 10.1023/A:1018456017643

Bleiweiss, R. (1999). Joint effects of feeding and breeding behaviour on trophic dimorphism in hummingbirds. Proc. R. Soc. B. 266: 2491–2497. 10.1098/rspb.1999.0951

Bonhomme, V., Picq, S., Gaucherel, C., and Claude, J. (2014). Momocs: Outline Analysis Using R. J. Stat. Softw. 56(13), 1–24. 10.18637/jss.v056.i13

Butler, M. A., Schoener, T. W., and Losos, J. B. (2000). The relationship between sexual size dimorphism and habitat use in Greater Antillean Anolis lizards. Evolution. 54(1), 259–272. 10.1111/j.0014-3820.2000.tb00026.x

Callieri, M., Ranzuglia, G., Dellepiane, M., Cignoni, P. and Scopigno, R. (2012). Meshlab as a complete open tool for the integration of photos and colour with high-resolution 3D geometry data. Comput. Appl. Q. Methods Archaeol. 406–416.

Chhaya, V., Reddy, S., Krishnan, A. (2023). Bill shape imposes biomechanical tradeoffs in cavity-excavating birds. Proc. R. Soc. B. 290: 20222395. 10.1098/rspb.2022.2395

Cook, R.D., Malkus, D.S., Plesha, M.E., Witt, R.J. (2002). Concepts and applications of finite element analysis. 4th ed. NY: Wiley & Sons.

Cooney, C.R., Bright, J.A., Capp, E.J.R., Chira, A.M., Hughes, E.C., Moody, C.J.A., Nouri, L. O., Varley, Z. K., and Thomas, G.H. (2017) Mega-evolutionary dynamics of the adaptive radiation of birds. Nature. 542, 344–347. 10.1038/nature21074

Dumont, E. R., Grosse, I. R., and Slater, G. J. (2009). Requirements for comparing the performance of finite element models of biological structures. J. Theor. Biol. 256(1), 96–103. 10.1016/j.jtbi.2008.08.017

Dumont, E. R., Samadevam, K., Grosse, I., Warsi, O. M., Baird, B., and Davalos, L. M. (2014). Selection for mechanical advantage underlies multiple cranial optima in new world leaf-nosed bats. Evolution. 68(5), 1436–1449. 10.1111/evo.12358

Emlen, D. J. (2008). The evolution of animal weapons. Annu. Rev. 39(1), 387–413. 10.1146/annurev.ecolsys.39.110707.173502

Fedorov, A., Beichel, R., Kalpathy-Cramer, J., Finet, J., Fillion-Robin, J.-C., Pujol, S., Bauer, C., Jennings, D., Fennessy, F., Sonka, M. et al. (2012). 3D Slicer as an image computing platform for the Quantitative Imaging Network. Magn. Reson. Imaging. 30, 1323–1341. 10.1016/j.mri.2012.05.001

Goodall, C. (1991). Procrustes methods in the statistical analysis of shape. J. R. Stat. Soc. Series B Stat. Methodol. 53, 285–339. 10.1111/j.2517-6161.1991.tb01825.x

Gunz, P., and Mitteroecker, P. (2013). Semilandmarks: a method for quantifying curves and surfaces. Hystrix. 24(1), 103–109. 10.4404/hystrix-24.1-6292

Habegger, M. L., Motta, P. J., Huber, D., Pulaski, D., Grosse, I. and Dumont, E. (2019). Feeding biomechanics in billfishes: investigating the role of the rostrum through finite element analysis. Anat. Rec. 303, 44–52. 10.1002/ar.24059

Harger, M., and Lyon, D. (1980). Further observations of lek behaviour of the Green Hermit Hummingbird Phaethornis guy at Monteverde, Costa Rica. Ibis. 122:525–530.

Hendry, A. P., Kelly, M. L., Kinnison, M. T., and Reznick, D. N. (2006). Parallel evolution of the sexes? Effects of predation and habitat features on the size and shape of wild guppies. J. Evol. Biol. 19(3), 741–754. 10.1111/j.1420-9101.2005.01061.x

Janicke, T. and Fromonteil, S. (2021). Sexual selection and sexual size dimorphism in animals. Biol. Lett.17: 20210251. 10.1098/rsbl.2021.0251

Klinkhamer, A. J., Woodley, N., Neenan, J. M., Parr, W. C. H., Clausen, P., Sánchez-Villagra, M. R., Sansalone, G., Lister, A. M. And Wroe, S. (2019) Head to head: the case for fighting behaviour in Megaloceros giganteus using finite-element analysis. Proc. R. Soc. B. 28620191873. 10.1098/rspb.2019.1873

Kodric-Brown, A., and Brown, J. H. (1978). Influence of economics, interspecific competition, and sexual dimorphism on territoriality of migrant rufous hummingbirds. Ecology. 59(2), 285–296. 10.2307/1936374

Lautenschlager, S. (2021). True colours or red herrings?: colourmaps for finite-element analysis in palaeontological studies to enhance interpretation and accessibility. R. Soc. Open Sci. 8 (11), 211357. 10.1098/rsos.211357

Leimberger, K.G., Dalsgaard, B., Tobias, J.A., Wolf, C. and Betts, M.G. (2022), The evolution, ecology, and conservation of hummingbirds and their interactions with flowering plants. Biol. Rev. 97: 923–959. 10.1111/brv.12828

MacDougall-Shackleton, E. and Harbison, H. (1998). Singing Behavior of Lekking Green Hermits. Condor. 100(1), 149–152. 10.2307/1369907

Marcé-Nogué, J., de Esteban-Trivigno, S., Escrig, C., and Gil, L. (2016). Accounting for differences in element size and homogeneity when comparing Finite Element models: Armadillos as a case study. *Palaeontol*. Electron. 19, 1–22. 10.26879/609

Marcé-Nogué, J. (2020). Mandibular biomechanics as a key factor to understand diet in mammals. In Mammalian Teeth—Form and Function (eds Martin, T. and Koenigswald, W. V.), pp. 54–80 (Verlag Dr. Friedrich Pfeil). 10.23788/mammteeth. 04

Marcé-Nogué J. (2022). One step further in biomechanical models in palaeontology: a nonlinear finite element analysis review. PeerJ 10:e13890 10.7717/peerj.13890

McCullough, E.L., Tobalske, B.W., and Emlen, D.J. (2014). Structural adaptations to diverse fighting styles in sexually selected weapons, Proc. Natl. Acad. Sci. U.S.A.111 (40) 14484–14488. 10.1073/pnas.1409585111

McGuire, J. A., Witt, C. C., Remsen, J. V., Jr., Corl, A., Rabosky, D. L., Altshuler, D. L., and Dudley, R. (2014). Molecular Phylogenetics and the Diversification of Hummingbirds. Curr. Biol. 24(8), 910–916. 10.1016/j.cub.2014.03.016

Medina, J., Irschick, D., Epperly, K., Cuban, D., Elting, R., Mansfield, L., Lee, N., Fernandes, A. M., Garzón-Agudelo, F., and Rico-Guevara, A. (2024). PicoCam: High-resolution 3D imaging of live animals and preserved specimens. Methods Ecol. Evol. 15, 1980–1989. 10.1111/2041-210X.14409

Menezes, J.C.T. and Palaoro, A.V. (2022). Flight hampers the evolution of weapons in birds. Ecol. Lett. 25, 624–634. 10.1111/ele.13964

Murphy, P. J., Rowe, A. J., Rayfield, E. J., and Janis, C. M. (2024). Finite element analysis of kangaroo astragali: A new angle on the ankle. J. Morphol. 285, e21707. 10.1002/jmor.21707

Perez, I. S., Bernal, V., and Gonzales, P. N. (2006). Differences between sliding semi-landmark methods in geometric morphometrics, with an application to human craniofacial and dental variation. J. Anat. 208, 769–784. 10.1111/j.1469-7580.2006.00576.x

R Core Team (2024). R: A Language and Environment for Statistical Computing. R Foundation for Statistical Computing, Vienna, Austria. <https://www.R-project.org/>.

Rayfield, E. J. (2007). Finite element analysis and understanding the biomechanics and evolution of living and fossil organisms. Annu. Rev. Earth Planet. Sci. 35(1), 541–576. 10.1146/annurev.earth.35.031306.140104

Rhodes, E. M., J. Borden and J. McCreadie. (2022). Quantification of physiological aging criteria utilizing window strike data. J. Field Ornithol. 93(4):12. 10.5751/JFO-00220-930412

Rico-Guevara, A., and Araya-Salas, M. (2015). Bills as daggers? A test for sexually dimorphic weapons in a lekking hummingbird. Behav. Ecol. 26(1), 21–29. 10.1093/beheco/aru182

Rico-Guevara, A., and Hurme, K. J. (2019). Intrasexually selected weapons. Biol. Rev. 94(1), 60–101. 10.1111/brv.12436

Rico-Guevara, A., Rubega, M. A., Hurme, K. J., and Dudley, R. (2019). Shifting Paradigms in the Mechanics of Nectar Extraction and Hummingbird Bill Morphology. Integr. Org. Biol. 1(1), oby006. 10.1093/iob/oby006

Rico-Guevara, A., Hurme, K. J., Rubega, M. A., and Cuban, D. (2023). Nectar feeding beyond the tongue: hummingbirds drink using phase-shifted bill opening, flexible tongue flaps and wringing at the tips. J. Exp. Biol. 226(Suppl_1), jeb245074. 10.1242/jeb.245074

Rico-Guevara, A., Sustaita, D., Hurme, K. J., Hanna, J. E., Jung, S., and Field, D. J. (2024). Upper bill bending as an adaptation for nectar feeding in hummingbirds. J. R. Soc. Interface. 2120240286. 10.1098/rsif.2024.0286

Rohlf, F. J. (1999). Shape statistics: Procrustes superimpositions and tangent spaces. J. Classif. 16, 197–223. 10.1007/s003579900054

Rowe, A. J., and Rayfield, E. J. (2022). The efficacy of computed tomography scanning versus surface scanning in 3D finite element analysis. PeerJ. 10, e13760. 10.7717/peerj.13760

Sansalone, G., Wroe, S., Coates, G., Attard, M. R., and Fruciano, C. (2024). Unexpectedly uneven distribution of functional trade-offs explains cranial morphological diversity in carnivores. Nat. Commun. 15(1), 3275. 10.1038/s41467-024-47620-x

Sargent, A. J., Groom, D. J. E., and Rico-Guevara, A. (2021). Locomotion and Energetics of Divergent Foraging Strategies in Hummingbirds: A Review. Integr. Org. Biol. 61(2), 736– 748. 10.1093/icb/icab124

Snow, B. K. (1973). The Behavior and Ecology of Hermit Hummingbirds in the Kanaku Mountains, Guyana. Wilson Bull. 85(2), 163–177. http://www.jstor.org/stable/4160317

Snow, B.K. (1974). Lek behaviour and breeding of Guy’s Hermit hummingbird *Phaethornis guy*. Ibis. 116: 278–297. 10.1111/j.1474-919X.1974.tb00125.x

Soons, J., Herrel, A., Genbrugge, A., Adriaens, D., Aerts, P., and Dirckx, J. (2012a). Multi-layered bird beaks: a finite-element approach towards the role of keratin in stress dissipation. J. R. Soc. Interface. 9, 1787–1796. 10.1098/rsif.2011.0910

Soons, J., Herrel, A., Aerts, P., and Dirckx, J. (2012b). Determination and validation of the elastic moduli of small and complex biological samples: bone and keratin in bird beaks. J. R. Soc. Interface. 9, 1381–1388. 10.1098/rsif.2011.0667

Stiles, F. G. (1980). The annual cycle in a tropical wet forest hummingbird community. Ibis. 122(3), 322–343. 10.1111/j.1474-919X.1980.tb00886.x

Stiles, F. G. (1995). Behavioral, ecological and morphological correlates of foraging for arthropods by the hummingbirds of a tropical wet forest. Condor. 97(4), 853–878. 10.2307/1369527

Stiles, F. G., and Wolf, L. L. (1979). Ecology and evolution of lek mating behavior in the long-tailed hermit hummingbird. Ornithol. Monogr. (27), iii–78.

Temeles E. J., Miller J S., and Rifkin J. L. (2010). Evolution of sexual dimorphism in bill size and shape of hermit hummingbirds (Phaethornithinae): a role for ecological causation. Phil. Trans. R. Soc. B. 365, 1053–1063. 10.1098/rstb.2009.0284

van der Meijden, A. and Kleinteich, T. (2017). A biomechanical view on stinger diversity in scorpions. J. Anat., 230: 497–509. 10.1111/joa.12582

Wainwright, S.A., Biggs, W.D., Currey, J.D., and Gosline, J.M. (1976). Mechanical design in organisms. Princeton, NJ: Princeton University Press.

Yanega, G., Pyle, P. and Geupel, G. (1997). The timing and reliability of bill corrugations for ageing hummingbirds.

Zhang, B., Baskota, B., Chabain, J. J., and Anderson, P. S. (2024). Curving expectations: the minimal impact of structural curvature in biological puncture mechanics. Sci. Adv. 10(33), eadp8157. 10.1126/sciadv.adp8157

